# Release of BEC-1 from CED-9 stimulates lysosome tubulation in *C. elegans* by engaging Class III PI3K complex I

**DOI:** 10.64898/2026.10.09.758037

**Authors:** Cristian A. Ricaurte-Perez, Aldrich K. Wilberg, Joshua P. Gill, Kathryn R. DeLeo, Olga Dubuisson, K. Adam Bohnert, Alyssa E. Johnson

## Abstract

The anti-apoptotic protein CED-9 and the autophagy protein BEC-1 have been linked to autophagy regulation, but how their interaction modulates this process is not fully understood. Here, we find that the physical dissociation of BEC-1 from CED-9 influences lysosomal morphology in addition to autophagic activity in *Caenorhabditis elegans*. We present evidence that when autophagy is stimulated via dietary restriction, BEC-1 is released from CED-9 and assembles with VPS-34 and other members of the PI3K complex I to promote PI3P enrichment at lysosomal membranes and subsequent membrane tubulation. Mutation of two hydrophobic amino acids within a predicted BH-3 like motif of BEC-1 is sufficient to disrupt its interaction with CED-9, thereby constitutively inducing lysosomal tubulation and increasing autophagic cargo turnover. This phenotype is dependent on VPS-34 activity, linking the CED-9/BEC-1 dissociation to PI3K-dependent lysosomal remodeling. Collectively, our work identifies the CED-9/BEC-1 interaction as a molecular switch that governs apoptosis, autophagy, and lysosomal morphology.

## Introduction

Dietary restriction (DR) remains one of the most reproducible approaches to extend lifespan across species, and multiple health benefits have been linked to DR in vertebrate and invertebrate organisms [1]. In humans, sustained DR for 6 months reduces plasma insulin levels and oxidative stress, and improves biomarkers associated with longevity [2]. Furthermore, long-term DR decreases the risk of atherosclerosis [3], reduces cardiometabolic risk factors [4], and improves liver function [5]. However, DR implementation in humans faces significant limitations, such as long-term adherence and some critical side effects, including loss of bone mineral density [6], accelerated fecundity decline [7–9] and mental health problems [10]. Thus, identifying the molecular and cellular mechanisms that mediate the beneficial effects of DR could lead to more practical therapeutic interventions to improve human health.

The autophagy/lysosome system has emerged as an attractive target to develop DR mimetics. Inhibiting autophagy negates DR-induced healthspan and lifespan extension in multiple species [11], suggesting autophagic activity is a central effector of DR’s pro-health and pro-longevity effects. Despite lysosomes having been canonically described as static, spherical-shaped organelles that break down cellular waste, it is now evident that lysosomes are much more dynamic and can change morphology in response to certain stimuli [12]. For example, we previously demonstrated in *Drosophila melanogaster* and *C. elegans* that vesicular lysosomes can morph into highly degradative tubular networks, which we refer to as tubular lysosomes (TLs) [13–15]. Moreover, we found that, in *C. elegans,* TLs are naturally stimulated by DR and can be experimentally induced by intestinal overexpression of *Drosophila SVIP (dSVIP)*, a well-characterized genetic stimulator of tubular lysosomes [15–18]. Significantly, forced lysosomal tubulation via *dSVIP* overexpression recapitulates the health benefits of DR, whereas inhibition of TL formation abolishes the positive effects of DR in *C. elegans* [15, 17]. The coordinated action of the insulin signaling pathway and mTOR pathway through their respective transcription factors, DAF-16/FOXO and HLH-30/TFEB, is indispensable to drive robust lysosomal tubulation under DR [19]. However, critical signaling events downstream of the DAF-16/HLH-30 regulatory axis that trigger membrane remodeling events necessary for the formation of TLs remains unclear.

In a previous RNAi-based genetic screen, we identified that VPS-34, a class III phosphatidylinositol-3-kinase (PI3K), and its scaffold subunit, BEC-1, are necessary for lysosomal tubulation in *C. elegans* [19], indicating that PI3K-dependent membrane signaling may play a major role in downstream mechanisms that link transcriptional programs to lysosomal remodeling. VPS-34 functions within two distinct complexes: PI3K complex I and PI3K complex II. These complexes share the same three core proteins (BEC-1, VPS-15, and VPS-34) but are functionally distinguished by their complex specific subunits; ATG-14/EPG-8 links complex I to autophagosome formation, whereas UVRAG directs complex II to endosomal trafficking [20]. VPS-34 is the catalytic kinase responsible for generating PI3P, a phosphoinositide that recruits downstream autophagy-related proteins containing PI3P-binding domains, including FYVE domains, to promote phagophore formation [21]. The remaining components contribute to the assembly, stability, and/or regulation of their respective complexes [20].

BEC-1 also binds to the anti-apoptotic protein CED-9 (*C. elegans* BCL-2 ortholog), and their physical association is thought to be a critical control point for apoptosis and autophagy activation [22]. Although the mechanism of this control is less clear in *C. elegans,* it has been shown that mammalian BECLIN-1 physically associates with BCL-2 via its BH3 domain, and it is thought that this association restrains BECLIN-1 from engaging PI3K complexes and thereby precludes autophagy activation [23–25]. This is supported by the observation that physically disrupting BECLIN-1/BCL-2 interaction triggers robust autophagy [26]. It has also been proposed that release of BECLIN-1 from BCL-2 may permit pro-apoptotic BH3-domain containing proteins to bind to BCL-2 and trigger release of downstream caspases [23, 27, 28]. Notably, specific disruption of BECLIN-1/BCL-2 interaction not only induces autophagy but is also sufficient to promote healthy aging in mice [26]. Whether BEC-1 dissociation from CED-9 would likewise activate autophagy and improve organismal aging in *C. elegans* remains unknown, as is the mechanism by which it would do so.

In this paper, we report that inhibition of any of the three core subunits of PI3K complexes I and II disrupts lysosomal tubulation, suggesting that proper TL assembly depends on the functionality and integrity of a core complex rather than relying on VPS-34 kinase activity alone. However, we demonstrate that complex I (containing EPG-8/ATG-14) specifically regulates lysosome tubulation, whereas complex II (containing UVRAG) is less critical to this specific process. Our data also indicate that the kinase activity of VPS-34 and lipid signaling is likely a critical event for lysosomal tubulation, since we observed an enrichment of PI3P in lysosomal membranes that correlated with membrane tubulations. Finally, we present evidence that BEC-1 dissociation from CED-9 also controls lysosomal tubulation. Specifically, abrogating BEC-1-CED-9 interaction via genetic mutation stimulates lysosomal tubulation constitutively and increases autophagic turnover, presumably by allowing BEC-1 to preferentially engage with PI3K complex I. Collectively, these findings indicate that the formation of tubular lysosomes during dietary restriction in *C. elegans* is regulated by PI3K complex I-dependent lipid signaling and requires BEC-1 to disengage from CED-9 for robust stimulation.

## Results

### Class III PI3K complex I regulates lysosome tubulation in *C. elegans*

Previously, we found that knockdown of *vps-34* or *bec-1* inhibited autophagic cargo turnover and the formation of tubular lysosomes in dietary-restricted *C. elegans* [19]. VPS-34 and BEC-1 both assemble and function in two distinct class III tetrameric complexes (complex I and complex II), but whether the other subunits of complex I and/or complex II are necessary for lysosome tubulation and enhanced cargo degradation in dietary-restricted *C. elegans* was unknown. To provide a more complete understanding, we screened the remaining PI3K complex subunits to determine if each complex is similarly required or if one complex plays a more predominant role in autophagic activity and lysosome tubulation. Complex I contains VPS-34, BEC-1, VPS-15, and EPG-8 (Figure 1A), while complex II contains VPS-34, BEC-1, VPS-15, and UVRAG (Figure 1B). In *C. elegans*, two predicted homologs of mammalian UVRAG have been identified: *Y34B4A.2* [29] and *T23G11.7* [30]. The latter encodes three predicted isoforms, T23G11.7a.1, T23G11.7a.2, and T23G11.7b.1 (Figure 1C). However, based on AlphaFold3 predictions, T23G11.7b.1 is structurally dissimilar to the other UVRAG homologs and is unlikely to interact with PI3K complex II (Figure 1D and Supplemental Figure 1A-C). Therefore, we focused our UVRAG analysis on *Y34B4A.2* and *T23G11.7* paralogs, which encode proteins that are most similar to human UVRAG.

**Figure 1:**
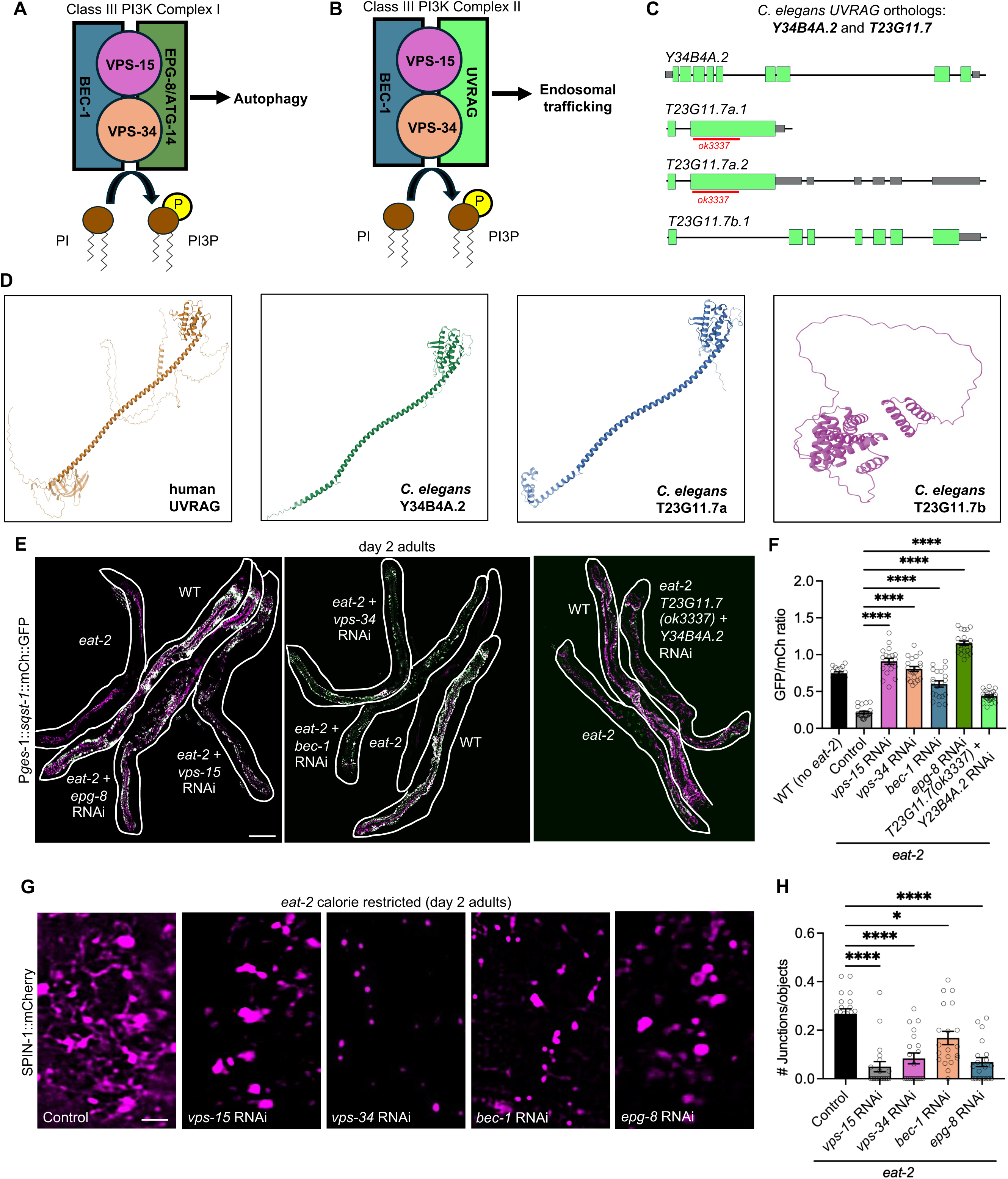
Class III PI3K complex I regulates lysosome tubulation. **(A)** Graphic model representing the class III PI3K complex I. **(B)** Graphic model representing the class III PI3K complex II. **(C)** Representation of *C. elegans UVRAG* orthologs highlighting the deletion region in the *T23G11.7 (ok3337)* mutant worms. **(D)** AlphaFold3 predictions of human UVRAG and *C. elegans* predicted UVRAG orthologs. **(E)** Gut-expressed SQST-1::mCherry::GFP in day 2 adults of WT (*N =* 17 worms)*, eat-2* mutants with L4440 control (*N =* 21 worms)*, eat-2* mutants with *vps-34* RNAi (*N =* 20 worms), *eat-2* mutants with *vps-15* RNAi (*N =* 20 worms), *eat-2* mutants with *bec-1* RNAi (*N =* 20 worms), *eat-2* mutants with *epg-8* RNAi (*N =* 21 worms), and *eat-2; T23G11.7 (ok3337)* double mutants with *Y34B4A.2* RNAi (*N* = 23 worms) (scale bar, 100μm). **(F)** GFP/mCherry ratios for the SQST-1::mCherry::GFP autophagy reporter in the genotypes indicated. ****p<0.0001. **(G)** Endogenously tagged SPIN-1::mCherry lysosome marker in calorie restricted *eat-2* day 2 worms with L4440 control, *vps-34* RNAi, *vps-15* RNAi, *bec-1* RNAi, and *epg-8* RNAi (*N =* 20 worms per condition) (scale bar, 5 μm). **(H)** Lysosomal junctions/objects with RNAi knockdown of class III PI3K complex I proteins in adult day 2 *eat-2* calorie-restricted worms (*p<0.05, ****p<0.0001). For all experiments, data were pooled from at least two independent experiments.

We first tested the necessity of each PI3K subunit for enhanced autophagic cargo turnover upon DR. Using a previously validated autophagy reporter, SQST-1::mCherry::GFP [15], we quantified autophagic turnover in *eat-2* dietary-restricted worms with each subunit inhibited by RNAi or via loss of function mutations. Consistent with our previous work, knockdown of either *vps-34* or *bec-1* via RNAi significantly blocked the increased autophagic cargo turnover normally observed in DR animals (Figure 1E-F). Likewise, knockdown of *vps-15* or complex I-specific *epg-8* by RNAi impaired the increased autophagic cargo degradation observed in *eat-2* mutants (Figure 1E-F). To simultaneously disrupt the two predicted *C. elegans UVRAG* homologs, we knocked down *Y34B4A.2* by RNAi in the *T23G11.7(ok3337)* mutant, which contains a deletion in exon 2 of both *T23G11.7a.1* and *T23G11.7a.2* isoforms (Figure 1C). Although disruption of the two UVRAG homologs slightly affected cargo turnover, this effect was substantially smaller than that observed following disruption of the PI3K complex I components (Figure 1E-F). Notably, the knockdown of *epg-8*, the complex I-specific subunit, produced the strongest block on autophagic turnover compared to any of the other subunits. Together, these results suggest that the increased degradation in autophagic substrates observed in dietary-restricted *C. elegans* depends predominantly on PI3K complex I activity, whereas PI3K complex II makes a relatively smaller contribution and does not fully compensate for the loss of the complex I during DR.

Previous work from our group demonstrated that autophagic cargo degradation in dietary-restricted worms relies on lysosome tubulation [15]. Thus, we next examined whether the individual components of PI3K complex I also contribute to lysosomal tubulation. Consistent with these findings, knockdown of each component of PI3K complex I produced a strong defect in lysosomal tubulation during DR (Figure 1G-H). These results indicate that PI3K complex I is not only required for the enhanced autophagic cargo turnover observed during DR but also plays an essential role to enable the formation of TLs.

### VPS-34-dependent PI3P lipids increase at lysosomal membranes in starved and dietary-restricted *C. elegans*

We observed that VPS-34 activity is necessary to induce lysosomal tubulation and the higher autophagic cargo turnover associated with TLs (Figure 1E-H). Because we also observed an enrichment of VPS-34 at tubular structures in starved worms [19], we hypothesized that PI3P enrichment at lysosomal membranes might facilitate expansion of membranes upon shifting from vesicles to tubules. To address this question, we used 2xFYVE, a PI3P-binding domain [31], fused to GFP as a quantitative reporter of PI3P-containing membranes. We assessed whether TLs triggered by nutrient-deprivation stimuli, such as starvation and DR, are enriched with 2xFYVE::GFP signal. Strikingly, when larval L1s were starved and tubular lysosomes activated, the distribution of the 2xFYVE::GFP signal transformed into structures that resembled the TL network (Figure 2A-B). Notably, lysosomes labeled with SPIN-1::mCherry showed significantly more co-localization with 2xFYVE::GFP after starvation compared to when animals were well-fed and lysosomes were predominantly vesicular (Figure 2A-B). In time-lapse imaging of starved animals, we observed dynamic 2xFYVE::GFP puncta merging with SPIN-1::mCherry-labeled lysosomes and subsequently diffusing along the lysosomal membrane lattice (Supplemental Video 1). Together, our data indicate that the relationship between PI3P and TLs during starvation might reflect active membrane remodeling associated with lysosomal tubulation.

**Figure 2:**
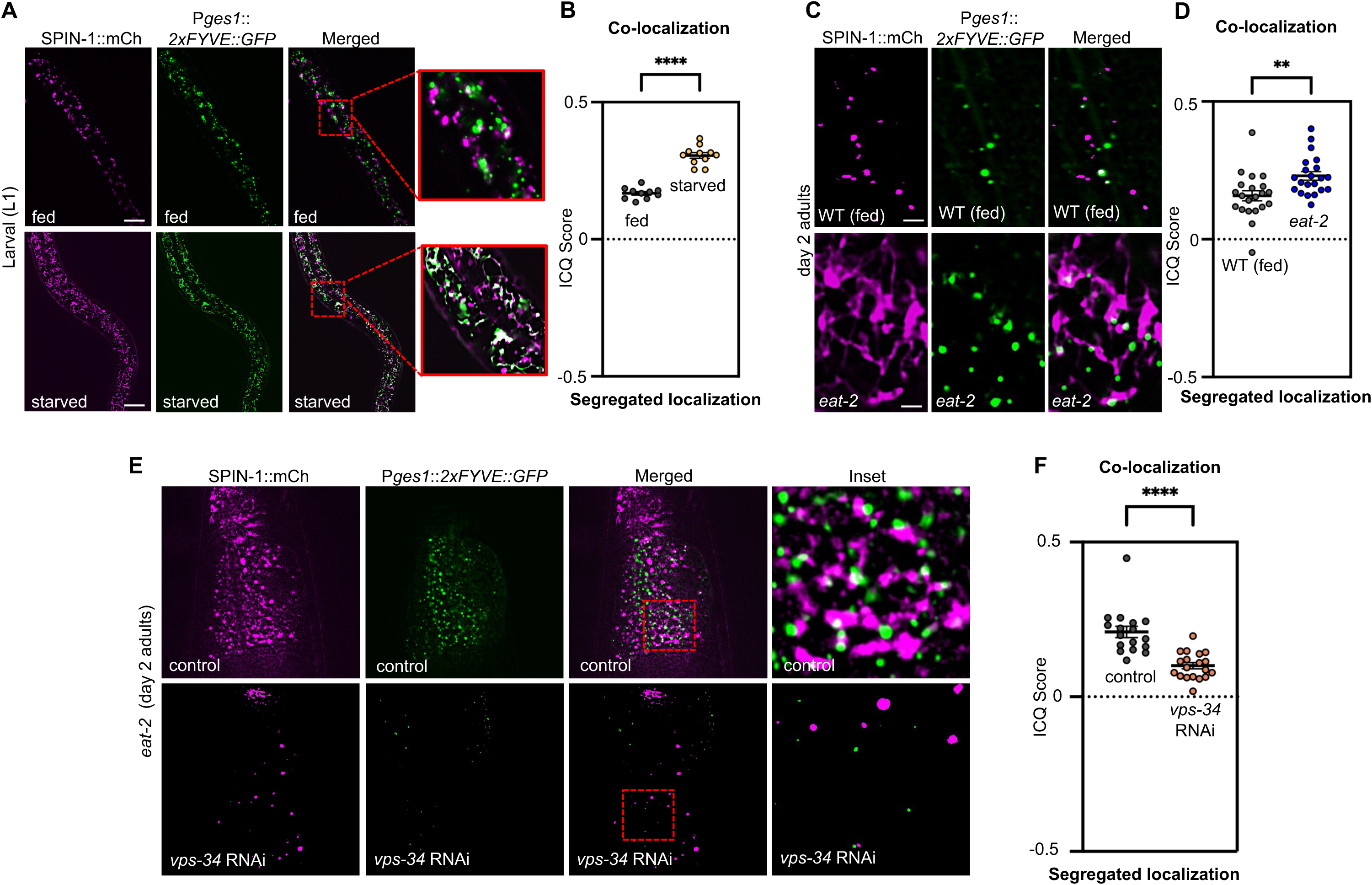
VPS-34-dependent PI3P lipids increase at lysosomal membranes in starved and dietary-restricted *C. elegans.* **(A)** Colocalization of endogenously tagged SPIN-1::mCherry lysosome marker with gut-expressed PI3P reporter, 2xFYVE::GFP, in fed and starved larval worms. **(B)** ICQ score quantified in fed and starved larval worms. Intensity correlation quotient (ICQ) values between 0 and 0.5 indicate colocalization and values between 0 and −0.5 indicate segregated fluorescence. Fed (*N* = 10 worms) and starved (*N* = 11 worms). Mean ± s.e.m. Unpaired two-tailed Student’s t-test (****p<0.0001). **(C)** Colocalization of endogenously tagged SPIN-1::mCherry lysosome marker with gut-expressed PI3P reporter, 2xFYVE::GFP, in wild-type and calorie-restricted *eat-2* mutant worms at day 2 of adulthood. **(D)** ICQ score quantified in wildtype and *eat-2* mutants. Intensity correlation quotient (ICQ) values between 0 and 0.5 indicate colocalization and values between 0 and −0.5 indicate segregated fluorescence. Wild-type control (*N* = 23 worms) and *eat-2* mutant (*N* = 22 worms). Mean ± s.e.m. Unpaired two-tailed Student’s t-test (**p<0.01) **(E)** Colocalization of endogenously tagged SPIN-1::mCherry lysosome marker with gut-expressed PI3P reporter, 2xFYVE::GFP, in calorie-restricted *eat-2* mutants on L4440 control and *vps-34* RNAi. **(F)** ICQ score quantified in *eat-2* mutants on control (*N* = 17 worms) and *vps-34* RNAi (*N* = 19 worms). Intensity correlation quotient (ICQ) values between 0 and 0.5 indicate colocalization and values between 0 and −0.5 indicate segregated fluorescence. Mean ± s.e.m. Unpaired two-tailed Student’s t-test (****p<0.0001). For all experiments, data were pooled from at least two independent experiments.

Next, we evaluated whether DR triggers a similar association between TLs and PI3P-positive structures. Although the distribution of the 2xFYVE::GFP signal did not change as dramatically as observed during starvation, we observed an increased overlap between SPIN-1::mCherry TLs and the 2xFYVE::GFP reporter in *eat-2* dietary restricted animals (Figure 2C-D). The distinct 2xFYVE::GFP signal distribution during starvation and DR may reflect differences in the severity of these nutrient-deprivation conditions. Starvation represents a more severe metabolic challenge and might induce a more extensive membrane remodeling need than DR.

Finally, we attempted to generate a VPS-34 kinase-dead mutant strain to evaluate if the observed phenotypes depend exclusively on the kinase activity of VPS-34 or if other VPS-34 functions are involved. However, our attempts to recover a homozygous VPS-34 kinase-dead mutant failed, consistent with an essential requirement for VPS-34 activity in viability [32]. As an alternative, we depleted *vps-34* by using RNAi. As expected, the colocalization between the mCherry signal marking TLs and the 2xFYVE::GFP signal was significantly reduced in dietary-restricted worms when *vps-34* was also knocked down (Figure 2E-F). We also observed a noticeable decrease in the total lysosomal membrane content with knockdown of *vps-34*, suggesting that PI3P production might be a general requirement for both lysosome biogenesis and tubulation. Collectively, our results demonstrate that TLs induced by nutrient deprivation are highly associated with PI3P-positive membranes, and that this association depends on the activity of VPS-34.

### Loss of *ced-9* promotes BEC-1 localization at lysosomal membranes and induces constitutive lysosome tubulation in well-fed animals

In PI3K complex I and II, BEC-1 acts as a scaffold and contributes to the assembly and localization of the other components of the complexes [33]. Before phagophore formation, mammalian BECLIN-1 has been reported to localize to intracellular compartments in the ER, where the complex assembles to generate PI3P [34]. Our data indicates that PI3P is enriched at the lysosomal membrane during lysosome tubulation (Figure 2) and that components of PI3K complex I are specifically required for TL formation (Figure 1). Thus, we wondered whether other components of complex I, namely BEC-1, would also redistribute to lysosomal membranes upon lysosome-tubulation events. To address this, we generated the reporter *Pges-1::bec-1::GFP* as an indicator of BEC-1 localization in the intestine and assessed if lysosomes labeled with SPIN-1::mCherry co-localized with BEC-1::GFP under fed and DR conditions. In well-fed animals, we observed that BEC-1::GFP displayed a sheet-like distribution that resembles the organization of the ER [35] (Figure 3A). Although we did not observe any significant variation in the distribution of BEC-1::GFP in fed or dietary-restricted animals, we observed significantly more colocalization between lysosomal signal with BEC-1::GFP signal in dietary-restricted animals compared to well-fed animals (Figure 3A-B). These data suggest that when autophagy is stimulated in DR conditions, BEC-1 engages with lysosomes more robustly, and this increased association might contribute to lysosome tubulation.

**Figure 3:**
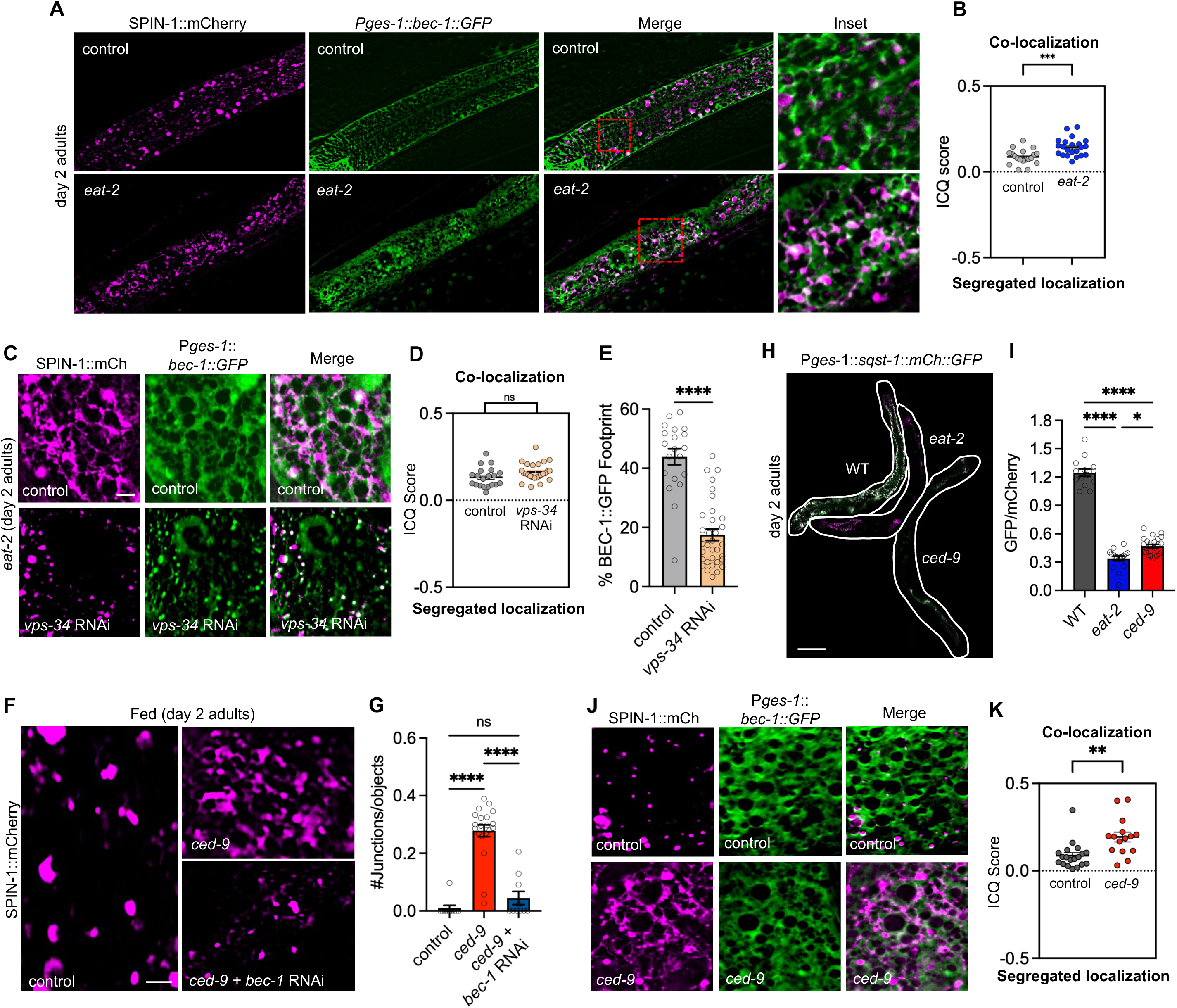
Loss of *ced-9* promotes BEC-1 localization at lysosomal membranes and induces constitutive lysosome tubulation in well-fed animals. (A) Colocalization of endogenously tagged SPIN-1::mCherry lysosome marker with gut-expressed *bec-1::GFP* in wildtype and calorie-restricted *eat-2* mutant worms. (B) ICQ score quantified in wildtype (*N* = 22 worms) and *eat-2* mutant (*N* = 19 worms) worms at day 2 of adulthood. Intensity correlation quotient (ICQ) values between 0 and 0.5 indicate colocalization and values between 0 and −0.5 indicate segregated fluorescence. Mean ± s.e.m. Unpaired two-tailed Student’s t-test (***p<0.001). (C) Colocalization of endogenously tagged SPIN-1::mCherry lysosome marker with gut-expressed *bec-1::GFP* in *eat-2* and *eat-2* + *vps-34* RNAi worms at day 2 of adulthood. **(D)** ICQ score quantified in *eat-2* mutant (*N* = 20 worms) and *eat-2* mutant + *vps-34* RNAi (*N* = 25 worms). Intensity correlation quotient (ICQ) values between 0 and 0.5 indicate colocalization and values between 0 and −0.5 indicate segregated fluorescence. Mean ± s.e.m. Unpaired two-tailed Student’s t-test (ns = not significant). **(E)** % BEC-1::GFP Footprint in gut expressed *bec-1::GFP* in *eat-2* (*N* = 20 worms) and *eat-2* + *vps-34* RNAi (*N* = 33 worms). Mean ± s.e.m. Unpaired two-tailed Student’s t-test (****p<0.0001). **(F)** Endogenously tagged SPIN-1::mCherry lysosome marker in wild-type, *ced-9* mutant, *ced-9* mutant *+ bec-1* RNAi. (Scale bar, 5 μm) **(G)** Number of junctions/objects with wildtype (*N =* 10 worms), *ced-9* mutant (*N =* 20 worms), *ced-9* mutant *+ bec-1* RNAi (*N =* 10 worms) (ns = not significant, ****p<0.0001) **(H)** Gut-expressed SQST-1::mCherry::GFP in wild-type, *eat-2* mutant, and *ced-9* mutant worms at day 2 of adulthood (scale bar, 100μm). **(I)** GFP/mCherry ratios for the SQST-1::mCherry::GFP autophagy reporter in wildtype (*N* = 11 worms), *eat-2* mutant (*N* = 17 worms), and *ced-9* mutant (*N* = 24 worms) worms at day 2 of adulthood (*p<0.05, ****p<0.0001). **(J)** Colocalization of endogenously tagged SPIN-1::mCherry lysosome marker with gut-expressed *bec-1::GFP* in wild-type and *ced-9* mutant worms at day 2 of adulthood. **(K)** ICQ score quantified in wildtype (*N* = 19 worms) and *ced-9* mutant (*N* = 15 worms) worms at day 2 of adulthood. Intensity correlation quotient (ICQ) values between 0 and 0.5 indicate colocalization and values between 0 and −0.5 indicate segregated fluorescence. Mean ± s.e.m. Unpaired two-tailed Student’s t-test (**p<0.01). For all experiments, data were pooled from at least two independent experiments.

Next, we evaluated if VPS-34 activity influences BEC-1 localization and its association with TLs under DR conditions. Interestingly, when we knocked down *vps-34* via RNAi, the sheet-like distribution of BEC-1::GFP dramatically changed to a punctuated pattern (Figure 3C-E). Surprisingly, we noticed that the degree of colocalization between BEC-1::GFP and lysosomes was similar regardless of lysosomal morphology. The punctuated BEC-1::GFP pattern colocalized with vesicular lysosomes to an extent comparable with BEC-1::GFP colocalization when TLs are deployed (Figure 3C–D). Potentially, this could suggest that lysosomal membranes get locked onto the BEC-1-associated ER membranes independently of VPS-34, but, without VPS-34 kinase activity, the membranes cannot remodel to form tubulated structures.

Besides its canonical role in regulating autophagy, BEC-1 also partially regulates apoptosis through its interaction with the anti-apoptotic protein CED-9 [22]. It is hypothesized that this physical association with CED-9 sequesters BEC-1 and limits the availability of BEC-1 for its autophagy-related functions. Our data demonstrates that BEC-1 and PI3K complex I are required for TL induction and enhanced autophagic cargo turnover. Thus, we sought to explore if loss of CED-9 would release BEC-1 and increase the pool of BEC-1 available to support autophagy and induce lysosomal tubulation. To test this, we first evaluated lysosomal morphology in well-fed animals with partial *ced-9* loss-of-function. Remarkably, lysosomal tubulation was robustly triggered in well-fed *ced-9* mutants (Figure 3F-G). These data suggest that a reduction in CED-9 activity is sufficient to induce robust TLs even in the absence of DR. To further determine whether this is a BEC-1-dependent phenotype, we knocked down *bec-1* via RNAi in *ced-9* mutants. Strikingly, we observed that a reduction of BEC-1 was sufficient to suppress the formation of TLs in well-fed *ced-9* mutants (Figure 3F-G), indicating that BEC-1 is necessary for the lysosomal tubulation induced by loss of CED-9. Consistently, *ced-9* mutants also had increased autophagic cargo degradation, reaching levels similar to those observed in dietary-restricted *C. elegans* (Figure 3H–I). Finally, we examined whether TLs induced in *ced-9* mutants also associated with the BEC-1::GFP reporter. Indeed, the TLs that formed in the *ced-9* mutant background showed increased co-localization with the BEC-1::GFP reporter (Figure 3J-K). Collectively, these findings support a model in which reduced CED-9 activity increases the availability of BEC-1 to boost autophagy and promote the formation of tubular lysosomes.

### Specific disruption of CED-9-BEC-1 interaction induces constitutive lysosome tubulation in well-fed animals and improves healthy aging

Our results indicate that CED-9 levels influence lysosomal morphology and that this effect depends on BEC-1 availability. However, CED-9 loss of function also triggers apoptosis [36], so we cannot rule out that the effects we were observing were due to increased BEC-1 availability or an effect of increased apoptosis. Thus, we next asked whether specifically disrupting the interaction between BEC-1 and CED-9, rather than altering the overall levels of either protein, would be sufficient to trigger TL formation.

In mammals, BECLIN-1 is composed of three major structural regions: the BH3 segment, a coiled-coiled domain, and a BARA domain. The BH-3 segment, which is approximately 20-25 amino acids in length, interacts with anti-apoptotic BCL-2 family proteins [24]. The coiled-coil domain mediates homodimerization and is essential to interact with the regulatory subunits ATG14 and UVRAG (EPG-8 and T23G11.7/Y34B4A.2 in *C. elegans*) [37]. The BARA domain has three structural repeats and contributes to membrane binding and regulation of complex I activity [38]. Since the interaction between BECLIN-1 and anti-apoptotic BCL-2 family proteins is mediated through the BH-3 domain, we focused on this region to identify critical residues that could impede the interaction between BEC-1 and CED-9 but retain the other functions of BEC-1. Although a specific BH3-like domain has not been characterized in *C. elegans* BEC-1, BH3 domains are known to have an amphipathic α-helical structure containing conserved hydrophobic residues that contribute to binding within the hydrophobic groove of BCL-2 family proteins [39]. Based on these features and some degree of sequence conservation with the mammalian BECLIN-1 BH3 region, we identified amino acids 33–53 of *C. elegans* BEC-1 as a candidate BH3-like region (Figure 4A). Within this region, we focused on the hydrophobic residues I38 and L42, which based on homology to mammalian BECLIN-1, are predicted to contribute to the interaction with CED-9 [39]. To specifically disrupt this interaction, we substituted isoleucine 38 (I38) and leucine 42 (L42) with alanine (Figure 4A). Importantly, AlphaFold3 structural predictions indicated that the substitution of these hydrophobic amino acids should be sufficient to disrupt the helix that directly interacts with the binding groove of CED-9 without affecting assembly of BEC-1 with PI3K complex I subunits (Supplementary Figure 2). To validate whether mutation of these two residues are sufficient to disrupt BEC-1/CED-9 binding, we assessed the interaction between these two proteins by co-immunoprecipitation. We used CRISPR to tag the N-terminus of CED-9 with 3xFLAG and the C-terminus of either wild-type BEC-1 or BEC-1(I38;L42A) with 2xHA to assess the interaction between these two proteins by co-immunoprecipitation. In wild-type conditions, BEC-1::2xHA co-immunoprecipitated with 3xFLAG::CED-9, confirming their association in *C. elegans* (Figure 4B). In contrast, we detected a significant reduction in the amount of BEC-1(I38;L42A)::2xHA co-immunoprecipitated with 3xFLAG::CED-9 (Figure 4B), indicating that mutations in the hydrophobic residues I38 and L42 contribute to the interaction between BEC-1 and CED-9, and their mutation is sufficient to abrogate BEC-1 interaction with CED-9. Importantly, we observed no significant changes to the total protein levels of either BEC-1 or CED-9 (Figure 4B). Thus, changes in these residues within the predicted BH-3 like motif are sufficient to abrogate the association with CED-9 without affecting the overall levels of either protein.

**Figure 4:**
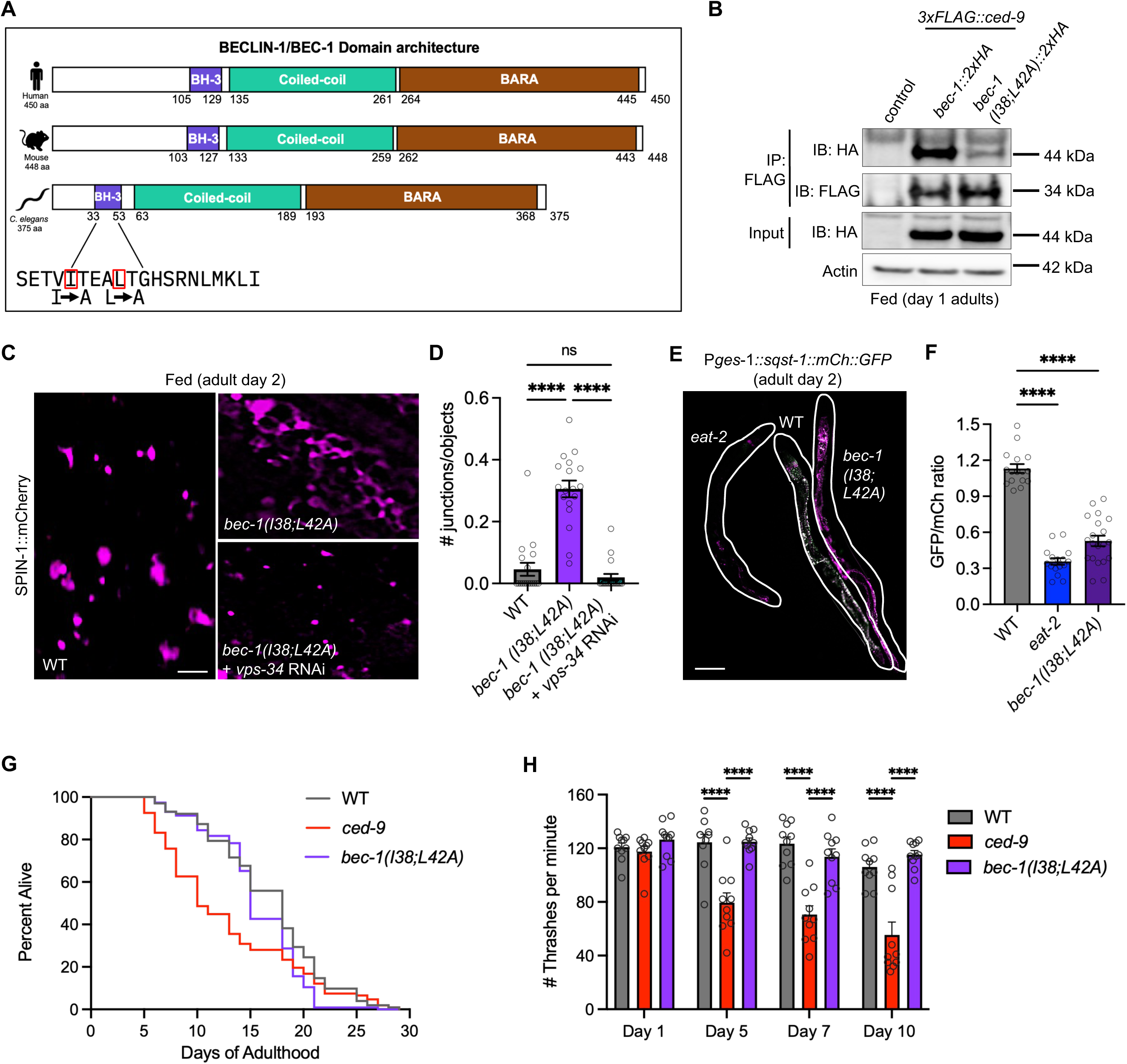
Specific disruption of CED-9-BEC-1 interaction induces constitutive lysosome tubulation in well-fed animals and improves healthy aging. (A) Representation of BECLIN-1/BEC-1 domain organization in humans, mice, and *C. elegans.* Proteins are shown from N-terminus to C-terminus left to right, with residue markers highlighting domain boundaries and total protein length. The BH-3 (BCL-2 homology 3), coiled-coil domain, and BARA domains are shown. The amino acid sequence for the *C. elegans* BH-3 with I38A and L42A substitutions that are present in the *bec-1(I38;L42A)* mutant worms are highlighted. (B) Co-immunoprecipitation of 3xFLAG::CED-9 with BEC-1::2xHA and BEC-1(I38;42A)::2xHA. (C) Endogenously tagged SPIN-1::mCherry lysosome marker in wildtype, *bec-1(I38;42A)*, and *bec-1(I38;42A)* + *vps-34* RNAi worms at day 2 of adulthood (scale bar, 5 μm). (D) Quantification of lysosomal junctions/objects in wildtype (*N =* 18 worms), *bec-1(I38;42A)* (*N =* 18 worms), and *bec-1(I38;42A)* + *vps-34* RNAi (*N =* 23 worms) worms at day 2 of adulthood. Mean ± s.e.m. One-way ANOVA test (ns = not significant, ****p<0.0001). Data were pooled from two independent experiments. (E) Gut-expressed *SQST-1::mCherry::GFP* in wildtype, *eat-2* mutant, and *bec-1(I38;42A)* worms at day 2 of adulthood (scale bar, 100μm). (F) GFP/mCherry ratios for the SQST-1::mCherry::GFP autophagy reporter in wildtype (*N =* 16 worms), *eat-2* mutant (*N =* 16 worms), and *bec-1(I38;42A)* (*N =* 20 worms) worms at day 2 of adulthood (****p<0.0001). Data were pooled from two independent experiments. (G) Lifespan analysis of wildtype (*N* = 102 worms), *ced-9 (N* = 107 worms), *bec-1(I38;42A)* (*N* = 115 worms). See source data file for descriptive statistics and log-rank test results. (H) Thrashing rates of the genotypes and ages indicated (*N* = 10 worms for all conditions and genotypes). Mean ± s.e.m. Two-way ANOVA with Šídák’s multiple comparisons test (\*\*\*\**p*<0.0001).

Next, we examined if the BEC-1(I38;L42A) substitution, like CED-9 loss of function, was sufficient to induce lysosomal tubulation even under well-fed conditions. Strikingly, we observed strong lysosomal tubulation in well-fed *bec-1(I38;L42A)* mutants compared to control animals (Figure 4C-D). As expected, this phenotype was suppressed upon the inhibition of VPS-34 activity (Figure 4C-D). Moreover, we observed improved cargo turnover in *bec-1(I38;L42A)* mutants, even to near similar levels detected in dietary-restricted *eat-2* mutants (Figure 4E-F). Collectively, our findings support a model in which breaking the physical interaction between CED-9 and BEC-1 increases the pool of BEC-1 available for autophagy-related functions and promotes lysosomal tubulation.

In previous studies, we have shown that constitutive induction of TLs in the intestine of *C. elegans* improves healthspan and healthy aging outcomes. Thus, we examined whether constitutive TL induction in *bec-1(I38;L42A)* mutants could also improve late-age health. Unlike *ced-9* loss of function mutants, which displayed a reduced lifespan compared to wild-type animals, likely due to the increased apoptotic activity, *bec-1(I38;L42A)* mutants did not display a reduction in lifespan (Figure 4G). Remarkably, however, *bec-1(I38;L42A)* mutants retained late-age mobility, while *ced-9* loss of function mutants displayed accelerated mobility decline with age compared to wild-type animals (Figure 4H). This is consistent with our previous observation that constitutive TL induction via *dSVIP* overexpression increased late-age mobility without affecting maximal lifespan [15]. Locomotor activity declines with age and is a strong indicator of late-age health in *C. elegans* [40]; thus, this improvement in late-age mobility indicates that disruption of BEC-1 and CED-9 interaction may extend healthspan by stimulating constitutive lysosome tubulation. Moreover, these data also suggest that mutation of critical residues in the BH3 domain of BEC-1 is sufficient to trigger autophagy without stimulating apoptosis.

## Discussion

The Class III PI3K, VPS-34, serves important roles in mediating diverse membrane remodeling events, including phagophore formation, vesicle sorting, and endocytic trafficking [41]. To regulate these diverse functions, VPS-34 assembles into two distinct lipid kinase complexes (complex I and complex II) that both contain BEC-1 and VPS-15 but differ in their inclusion of either ATG-14/EPG-8 or UVRAG [42]. In this study, we showed that PI3K complex I, rather than PI3K complex II, is the primary regulator of lysosomal tubulation and enhanced autophagic cargo turnover in dietary-restricted *C. elegans*. This is supported by our data that depleting EPG-8, the complex I-specific subunit, strongly disrupted autophagy cargo turnover and lysosome tubulation, while depleting *C. elegans* UVRAG orthologs had a much milder effect on cargo turnover. These data are also consistent with previous work demonstrating that PI3K complex I functions more specifically in autophagy, while complex II functions predominantly in endocytic trafficking [20]. Our work now adds lysosomal tubulation as another autophagy-related function regulated by PI3K complex I.

The requirement for VPS-34 in lysosomal tubulation hints at a potential role for PI3P production in priming lysosomal membranes for tubulation. Accordingly, we found PI3P-positive membranes become enriched at lysosomal membranes upon the morphological shift from vesicles to tubules, suggesting that PI3K complex I activity may be a prerequisite for lysosomal tubulation. How PI3P lipid modification promotes lysosomal tubulation remains to be determined; however, it is likely that increased localized enrichment of PI3P at lysosomal membranes promotes recruitment of membrane remodeling factors that contain PI3P-specific binding domains, such as FYVE and PX domains [43]. Potentially, factors that could promote membrane bending and extrusion (e.g. BAR domain-containing proteins), proteins that can promote lipid delivery, and/or proteins that promote homotypic membrane fusion could be recruited to lysosomal membranes via local enrichment of PI3P and subsequently cooperate to drive membrane tubulation [44]. Additionally, in previous work we found that TLs are highly dynamic and require the microtubule cytoskeleton for robust tubulation events [13]. Thus, PI3P lipid accumulation could also help recruit proteins that couple lysosomal membranes to the microtubule cytoskeleton or motor proteins to promote membrane dynamics. Identification of the key lipid remodeling factors that are recruited to lysosomes to promote tubulation will be an important next step.

Finally, we present evidence that dissociation of BEC-1 from the anti-apoptotic protein CED-9 is a major regulatory step that influences the ability of BEC-1 to participate in autophagy-related functions, including lysosome remodeling. Most notably, specific disruption of BEC-1 binding to CED-9 via mutation of the BH3 domain in BEC-1 was sufficient to drive lysosome tubulation constitutively and increase autophagic cargo turnover. Moreover, unlike *ced-9* loss of function mutants that showed reduced lifespan, specific mutation of the BEC-1 BH3 domain did not cause reduced lifespan but in fact improved late-age health. These data are consistent with previous studies demonstrating that targeted disruption of mammalian BECLIN-1 binding to BCL-2 heightens autophagic activity and extends lifespan in mice [26]. Thus, mutation of the BH3 domain could provide a useful means of activating BEC-1-dependent autophagy functions without apoptosis activation. Indeed, compounds that selectively disrupt BECLIN-1 binding to BCL-2 have become an emerging approach for geroprotective therapies [45, 46].

Collectively, our study supports a model in which under non-autophagic conditions, CED-9 sequesters BEC-1 and restrains its ability to engage in autophagic and lysosome remodeling functions. This model mirrors the established role of mammalian BECLIN-1, in which binding of BECLIN-1 to anti-apoptotic BCL-2 family proteins through its BH3 domain limits BECLIN-1-dependent autophagy [23, 24]. Our results indicate that, once freed from CED-9, BEC-1 amplifies autophagic turnover in part via TL stimulation. This could explain the improved healthspan observed in mice [23], and now *C. elegans*, upon chronic dissociation of BECLIN-1/BEC-1 from BCL-2/CED-9. Thus, the interaction between CED-9 and BEC-1 serves as a key regulatory point that links apoptosis, autophagy, and lysosomal morphology, and this mechanism of control appears to be broadly relevant to multiple animal species.

## Materials and Methods

### Strain generation

Supplementary Table 1 provides a complete list of strains used in this study. All strains used in this study were generated using standard genetic crosses and microinjection [47]. For genetic crosses, transgenes expressing fluorescent proteins were tracked by stereomicroscopy, and gene deletions and mutations were verified by PCR and/or sequencing. For microinjection, constructs were injected individually or in combination into the gonad of adult hermaphrodites, each at a concentration of 25 ng/µl.

#### Pges-1::2xFYVE::GFP::unc-54 3′ UTR

To generate *Pges-1::2xFYVE::GFP::unc-54 3′ UTR*, the 2xFYVE::GFP sequence was PCR-amplified from *ced-1*p::2xFYVE::GFP transgenic *C. elegans* genomic DNA using primers containing Gateway attB sites upstream of the start codon: Forward: 5′-GGGG ACA AGT TTG TAC AAA AAA GCA GGC TCA AAA GACTCTAGAGGATCCATGG-3′ and Reverse: 5′-GGGG AC CAC TTT GTA CAA GAA AGC TGG GTCTATTTGTATAGTTCATCCATGCCATG-3′. The PCR product was cloned into pDONR221 using BP clonase (ThermoFisher), and the insert sequence was verified. The resulting pDONR221-*2xFYVE::GFP* plasmid was then combined with pDONR P4-P1r P*ges-1* and pDONR P2R-P3 *unc-54* 3′ UTR in a pDEST R4-R3 Gateway destination vector using LR clonase (ThermoFisher) to generate the final expression construct.

#### Pges-1::bec-1::GFP::unc-54 3′ UTR

To generate *Pges-1::bec-1::GFP::unc-54 3′ UTR*, the *bec-1* genomic sequence excluding the endogenous stop codon was PCR amplified from N2 *C. elegans* using primers containing flanking Gateway attB sites; additionally, a XbaI restriction site was added to the Reverse primer in between the end of the *bec-1* annealing region and the attB site. Forward: 5’-GGGGACAAGTTTGTACAAAAAAGCAGGCTCAAAAATGACGACCCAACGAAGCC-3’ and Reverse: 5’-GGGGACCACTTTGTACAAGAAAGCTGGGTTCTAGAAATAGGCGATCTGAGAGCATCGC-3’. The *bec-1* genomic PCR product was cloned into pDONR221 using BP clonase (ThermoFisher), and the insert sequence was verified by Sanger sequencing. Subsequently, a Gibson reaction was performed to clone in the GFP coding sequence in frame with *bec-1.* Briefly, the pDONR221 *bec-1* plasmid was digested and linearized using XbaI and extracted from a DNA agarose gel. Then, the GFP coding sequence was amplified from a previously made pDONR221 *GFP* plasmid using primers containing 35 base pairs of homology to the pDONR221 *bec-1* plasmid; the linearized pDONR *bec-1* plasmid and GFP PCR product were used to perform the Gibson reaction. Forward: 5’-GCTCAATGCGATGCTCTCAGATCGCCTATTTCTAGAATGAGTAAAGGAGAAGAACTTTTC-3’ and Reverse: 5’ GCCAACTTTGTACAAGAAAGCTGGGTTCTAGA CTATTTGTATAGTTCATCCATGCCATG-3’. The *GFP* coding sequence was verified by Sanger sequencing. The resulting pDONR221 *bec-1::GFP* plasmid was then combined with pDONR P4-P1r P*ges-1* and pDONR P2R-P3 *unc-54* 3′ UTR in a pDEST R4-R3 Gateway destination vector using LR clonase (ThermoFisher) to generate the final expression construct. After injection, the extrachromosomal array was integrated into the endogenous genome by exposing 30 L4 worms to 30 mJ/cm² of ultraviolet radiation in a Stratalink UV crosslinker on a plate with no OP50 bacteria. Worms were transferred to plate with OP50 bacteria to recover and produce progeny. F2 worms were singled and homozygous integrants were selected based on 100% fluorescent progeny in the F3 generation. The resulting integrated strain was then outcrossed 6x to N2 WT worms before use.

### CRISPR-Cas9 Modified strains

The following *C. elegans* edited alleles were generated using a CRISPR/Cas9-mediated genome-editing strategy. The method for the generation of each strain is detailed below.

#### bec-1(lou30[I38A, L42A]) strain

The BH-3-like domain of BEC-1 was mutated by substituting isoleucine 38 (I38) and leucine 42 (L42) to alanine. The *bec-1*-targeting crRNA (5′-CAUCCGAGAUGAGCUUCAUGGUUUUAGAGCUAUGCU-3′) was designed and synthesized by Integrated DNA Technologies (IDT). The ssODN donor template 5′ GCGCCTTTAAGAATTTATTTTAAA AATTGATCCCAAAAATGTGTTTTTTACATAATATCAATTTTTAGAGCGAAACAGTTGCC ACAGAAGCTGCCACTGGACATTCTCGAAATCTTATGAAGCTCATCTCGGATGCTCAAT TCCCATCTGATGCTCCAGTTTGTAATGTTGGTTTTGA-3′ was designed to introduce the I38A and L42A substitutions and contained an 80bp upstream homology arm and a 60bp downstream homology arm flanking the edited region. An additional silent mutation was incorporated into the repair template to disrupt the endogenous BspHI restriction site located near the mutation region. As co-marker, we used a *dpy-10* crRNA 5 - GCUACCAUAGGCACCACGAGGUUUUAGAGCUAUGCU-3′, and the *dpy-10* ssODN 5′-ACTTGAACTTCAATACGGCAAGAT GAGAATGACTGGAAACCGTACCGCATGCGGTGCCTATGGTAGCGGAGCTTCACATGG CTTCAGACCAACAGCCTAT-3′ as previously described [48]. CRISPR/Cas9 injection mix was prepared in 200 mM HEPES (pH 7.4) and 1 mM KCl with final concentrations of 4 μg/μl tracrRNA, 8 μg/μl dpy-10 crRNA, 8 μg/μl target crRNA, 500 ng/μl *dpy-10* ssODN, 500 ng/μl target ssODN and 10 μg/μl Cas9 (Integrated DNA Technologies; catalog no. 1081059) and injected into the germline of N2 wild-type worms.

F1 animals displaying the co-CRISPR *dpy-10* mutant phenotype were singled and genotyped for the desired *bec-1(I38;L42A)* mutation. To genotype the mutation, the *bec-1* target sequence was PCR amplified using the primers: forward: 5′ GCAAAAAGTTTATGGTAGTC-3′; and reverse, 5′-CGAATACTTGTCTGGAAATTC-3′, which anneal outside the homology arms. The PCR products were subjected to a restriction enzyme digest with the BspH1 enzyme for 1hr at 37C before resolving on an agarose gel. Homozygous WT animals yielded 2 products of approximately 480 and 900bps. Homozygous *bec-1(I38, L42A)* yielded a single product of approximately 1380bp and heterozygous animals produced three band products of approximately 480, 900, and 1380bps. Correct editing of the *bec-1* locus was confirmed by sequencing. We also confirmed no additional mutations were incorporated during the CRISPR engineering. A single founder homozygous for *bec-1*(*lou30[I38A, L42A])* and carrying the *dpy-10* co-CRISPR allele was identified. The founder was outcrossed to N2 wild-type animals to eliminate the *dpy-10* co-CRISPR allele. Progeny were subsequently screened to confirm the presence of the *bec-1(I38A, L42A)* mutation and loss of the *dpy-10* phenotype.

#### bec-1(lou10[bec-1::2xHA]) strain

A 2xHA tag was introduced at the endogenous C-terminus of *bec-1* using the CRISPR/Cas9 co-CRISPR strategy described above. The *bec-1* targeting crRNA sequence was 5′-ATCCCGTGAAATCACTAAAT-3′. A linear dsDNA repair template encoding a Glycine-Serine linker followed by the 2xHA tag was generated by PCR amplification of the linker::2xHA cassette from plasmid pHD-2xHA-ScarlessDsRed (Addgene #8082) using primers containing 65bp homology arms as 5′ tails matching sequences flanking the *bec-1* stop codon. The primer sequences were: forward, 5′-CAATGGCTCTAATGCTCACAGATCTCAAAGCTGCGTGTGCTCAATGCGATGCTCTCAGATCGCCTATTGGTGGATCTGGAGGTTCCGGCG-3′; and reverse, 5′-AAATTGAAAAAAAAATTTCTAGCGTTGAAAATGTTATTCTACATTTGATCCCGTGAAATCACTAAATAGCATAATCTGGAACATCATATG-3′. The resulting 231 bp PCR product, containing the upstream homology arm, linker::2xHA coding sequence, and downstream homology arm, served as the repair template.

F1 animals displaying the co-CRISPR *dpy-10* mutant phenotype were selected and genotyped for *bec-1::2xHA* insertion. The target locus was amplified by PCR using the primers: forward, 5′-CATGCTGATTCACTGGTTGACAATGGTG-3′ and reverse: 5′-GCCTTTTCCCCAAACTTGTACCATATG-3′ designed to flank the edited region. Homozygous wild-type animals lacking the 2xHA tag yielded a 320bp PCR product, whereas animals homozygous for *bec-1::2xHA* yielded a 420 bp product. A single founder homozygous for *bec-1::2xHA* and carrying the *dpy-10* co-CRISPR allele was identified. The founder was outcrossed to N2 wild-type animals to outcross the *dpy-10* co-CRISPR allele. Progeny were subsequently screened to confirm the presence of the *bec-1::2xHA* allele and loss of the *dpy-10* mutation.

#### ced-9(lou9[3xFLAG::ced-9]) strain

A 3xFLAG tag was introduced into the endogenous N-terminus of *ced-9* using the CRISPR/Cas9 co-CRISPR strategy described above. The *ced-9* targeting crRNA sequence was 5′-TTAGATGACACGCTGCACGG-3′. A linear dsDNA repair template encoding the 3xFLAG tag was generated by PCR amplification of the 3xFLAG cassette from plasmid pHD-3xFLAG-ScarlessDsRed (Addgene #80820) using primers containing 60bp homology arms as 5′ tails matching sequences flanking the *ced-9* insertion site. The primer sequences were: forward: 5′-CTCTTCGTTTTTTTTTCTCTGTCTAGAATGTATATTATGATTATGAAAACGAATAAAATTTTAGATGGACTACAAAGACCATGACGGTG-3′; and reverse: 5′-CGCCATCGTTCGTCGCCGATACGCCGGATTCGTCAGCGAGTTGTCCGCCGTGCAGCGTGTCATGCCCTTGTCATCGTCATCCTTGTAATC-3′. The resulting 200 bp PCR product, containing the upstream homology arm, 3xFLAG coding sequence, and downstream homology arm, served as the repair template.

F1 animals displaying the co-CRISPR *dpy-10* mutant phenotype were selected and genotyped for the desired *3xFLAG::ced-9* insertion. The target locus was amplified by PCR using primers designed to flank the edited region (forward, 5′-CAGCCCTCGACAATTCTCGTCAATTTAC-3′; reverse, 5′-GTCCTGAGCATCACTATTGATTCC-3′). Homozygous wild-type animals lacking the 3xFLAG tag yielded an approximately 280 bp PCR product, whereas animals homozygous for the *3xFLAG::ced-9* allele yielded an approximately 350bp product. A single founder homozygous for *3xFLAG::ced-9* and carrying the *dpy-10* co-CRISPR allele was identified. The founder was outcrossed to wild-type animals to eliminate the *dpy-10* co-CRISPR allele. Progeny were subsequently screened to confirm the presence of the *3xFLAG::ced-9* allele and loss of the *dpy-10* mutation.

#### bec-1(lou31[I38A, L42A]::2xHA) strain

The *bec-1(lou31[I38A, L42A]::2xHA)* strain was generated by introducing the 2xHA tag into the previously generated *bec-1(lou30[I38A, L42A])* background using the CRISPR/Cas9 co-CRISPR strategy described above. The *bec-1(lou30[I38A, L42A])* strain was used as the parental strain for CRISPR/Cas9-mediated insertion of the 2xHA tag. F1 animals displaying the co-CRISPR *dpy-10* mutant phenotype were screened for the *bec-1::2xHA* insertion using the genotyping strategy described above. The missense mutations in the final strain were re-confirmed by sequencing. The *dpy-10* co-CRISPR allele was subsequently removed by outcrossing to wild-type animals.

### Animal maintenance

Worms were raised at 20°C on NGM agar (51.3 mM NaCl, 0.25% peptone, 1.7% agar, 1 mM CaCl2, 1 mM MgSO4, 25 mM KPO4, 12.9 µM cholesterol, pH 6.0). Fed worms were maintained on NGM agar plates previously seeded with *E. coli* OP50 bacteria. Synchronous populations of worms were obtained by bleaching gravid hermaphrodites. Briefly, gravid worms were vortexed in 1 mL bleaching solution (0.5 M NaOH, 20% bleach) for 5 min to isolate eggs, and eggs were then washed three times in M9 buffer (22 mM KH2PO4, 42 mM Na2HPO4, 85.5 mM NaCl, 1 mM MgSO4) before plating. To obtain starved L1 animals, bleached eggs were spotted on NGM agar that lacked OP50 bacteria, and plates were maintained at 20°C for 24–48 h before imaging. For aging experiments, synchronous populations of animals were established by bleaching gravid worms. In all aging experiments, adult worms were picked onto fresh OP50-seeded NGM plates every day to separate adults from their progeny until reproductive cessation.

### RNAi experiments

The *vps-34*, *vps-15*, *bec-1,* and *epg-8* RNAi clones were obtained from the Julie Ahringer RNAi collection [49] and verified by DNA sequencing. For RNAi experiments, synchronous populations of animals were grown on RNAi plates (NGM plus 100 ng/µl carbenicillin and 1 mM IPTG) that had been seeded with bacteria expressing the RNAi clone. An empty L4440 vector was used as a negative control. The *Y34B4A.2* RNAi clone was generated from *C. elegans* cDNA. The *Y34B4A.2* was amplified using the forward primer: 5’-CGGAGGATGTGGCTGTGGATAGTGA-3’ and reverse primer: 5’-GCTGCAGTTGTTTCCCGTTCG-3’. The resulting PCR product was cloned into an RNAi feeding vector using Gateway BP and LR recombination reactions [50]. The resulting plasmid was transformed into *E. coli,* and positive clones were confirmed by sequencing.

### Lifespan analysis

Synchronous populations of worms were transferred as late L4s to NGM plates seeded with OP50 bacteria. Animals that exploded, bagged, or crawled off plates were censored during analysis as is standard practice. Lifespans were analyzed using OASIS 2 software [51], and statistical significance was assessed using a log-rank test. Each lifespan plot represents a single independent biological replicate.

### Thrashing assay

Synchronous populations of worms were transferred as late L4s to NGM plates seeded with OP50 bacteria. Worms were transferred to fresh plates every day to separate adults from their progeny. To score thrashing rates, individual worms were transferred into a drop of M9 buffer on an NGM plate, and the number of body thrashes were counted in a 1 min period. Each worm represents an independent biological replicate.

### Microscopy

For *C. elegans* whole animal imaging, 4% agarose (Fisher Bioreagents) pads were dried on a Kimwipe (Kimtech) and then placed on top of a Gold SealTM glass microscope slide (ThermoFisher Scientific). A small volume of 10mM levamisole (Acros Organics) was spotted on the agarose pad. Worms were transferred to the levamisole spot, and a glass cover slip (Fisher Scientific) was placed on top to complete the mounting.

### Western blot and co-immunoprecipitation

For each genotype, day 1 adult worms were washed off 100mm plate in M9 buffer and centrifuged briefly to pellet worms. For co-IPs, worms from four 100mm plates were pooled for each sample. Worms were lysed using a motorized grinder in RIPA buffer (sc-24948, Santa Cruz Biotechnology) supplemented with halt phosphatase inhibitor cocktail (78420 Thermo Fisher) to extract total protein, and the tissue was centrifuged to pellet the solid debris. For co-immunoprecipitation, the soluble lysate was collected and incubated with 25µl of magnetic ANTI-FLAG M2 beads (M8823, Millipore) overnight at 4°C with gentle mixing. Beads were subsequently washed four times with NP-40 buffer. Following the final wash, 5X sample buffer (1M Tris-HCl, pH 6.8, 1 mM DTT, 20% SDS, 60% glycerol, and bromophenol blue) was added to the beads, and samples were boiled for 10min. An aliquot of the lysate collected before immunoprecipitation was retained as the input sample. Proteins were resolved by SDS-PAGE on a 4-12% Bis-Tris gel (Invitrogen, Thermo Fisher Scientific), subsequently transferred to a nitrocellulose membrane, and blocked in 4% milk in Tween-containing buffer for 1 hour at room temperature. For immunoblotting, membranes were incubated overnight at 4°C with anti-HA tag antibody (1:1000; ab9110, Abcam) or monoclonal ANTI-FLAG M2 antibody (1:1000; F3165, Millipore) diluted in Tween-containing buffer. Subsequently, membranes were washed three times in PBS containing 0.1% Tween 20 and incubated for 1 hour at room temperature with HRP-conjugated goat anti-rabbit IgG or goat anti-mouse IgG secondary antibodies diluted in 1% milk in Tween-containing buffer. Proteins were detected using ECL chemiluminescent reagent (170-5060, Bio-Rad) and imaged using a Bio-Rad GelDoc imager.

### AlphaFold structural prediction

AlphaFold Server powered by AlphaFold3 [52] was used to generate structural predictions of full length human UVRAG, *C. elegans* Y43B4A.2, and *C. elegans* T23G11.7a and T23G11.7b isoforms. Additionally, AlphaFold Server was used to predict the structure of the *C. elegans* PI3K complex II composed of BEC-1, VPS-15, VPS-34, and each of the UVRAG orthologs, Y43B4A.2, T23G11.7a, or T23G11.7b. To investigate the potential effects of mutations within the BEC-1 BH3 domain, structural models were generated for complexes containing wild-type BEC-1 or BEC-1 (I38A, L42A) mutant in complex with VPS-34, VPS-15, and EPG-8.

### Image analysis

Images were processed using LAS X software (Leica) and Fiji/ImageJ (NIH). Lysosome networks were analyzed using “Skeleton” analysis plugins in Fiji. Briefly, images were converted to binary 8-bit images and then to skeleton images using the “Skeletonize” plugin. Skeleton images were then quantified using the “Analyze Skeleton” plugin. Number of objects, number of junctions, and object lengths were scored. An “object” is defined by the Analyze Skeleton plugin as a branch connecting two endpoints, an endpoint and junction, or two junctions. Junctions/object was used as a parameter to quantify network integrity. For analyzing fluorescence intensity, the gut tissue was outlined using the free-draw tool in Fiji/ImageJ, and average fluorescence intensity of the outlined area was measured. For SQST-1::mCherry::GFP fluorescence ratio experiments, 50% laser intensity, 300ms exposure time, and 100% Fluorescence Intensity Manager settings were used. For co-localization analysis, whole worm images were analyzed using Fiji’s colocalization plugin Coloc2 to determine the colocalization of green and red fluorescence, which was reported as the intensity correlation quotient (ICQ) score [53]. ICQ values that tend toward 0 indicate random fluorescence patterns, segregated localization values will tend toward −0.5, and values for colocalization of the fluorescent signals will tend towards 0.5 [54, 55].

## Statistical analyses

Data were statistically analyzed using GraphPad Prism. For two sample comparisons, an unpaired t-test was used to determine significance (α = 0.05). For three or more samples, a one-way ANOVA with Dunnett’s, Tukey’s, or Šídák’s multiple comparisons was used to determine significance (α = 0.05). For grouped comparisons, a two-way ANOVA with Šídák’s multiple comparisons was used to determine significance (α = 0.05). Statistical significance of lifespan data was determined using a log-rank test.

## Data Availability

All data are available in the main text or the supplementary materials. Additional information on data sources is available upon request from the corresponding author. All unique materials used in the study are available from the authors or from commercially available sources.

## Acknowledgments

The authors thank all members of the Bohnert and Johnson labs for helpful discussions on this project. Funding for this project comes from: NIH-NIA grant RF1AG090648 (AEJ and KAB), NIH-NIGMS grant R35GM138116 (AEJ), and an American Heart Association predoctoral fellowship (CRP).

## Author contributions

Conceptualization: CRP, AKW, JPG, KAB, AEJ; Methodology: CRP, AKW, JPG, KRD, OD; Formal Analysis: CRP, AKW; Investigation: CRP, AKW, OD; Resources: CRP, JPG, KRD, OD; Data Curation: CRP, AKW, KRD; Visualization: CRP, AKW; Funding acquisition: CRP, KAB, AEJ; Project administration: CRP, AEJ; Supervision: KAB, AEJ; Writing – original draft: CRP, AEJ; Writing – review & editing: CRP, AKW, JPG, KRD, KAB, AEJ

