## Supplemental Figures for "Release of BEC-1 from CED-9 stimulates lysosome tubulation in *C. elegans* by engaging Class III PI3K complex I"

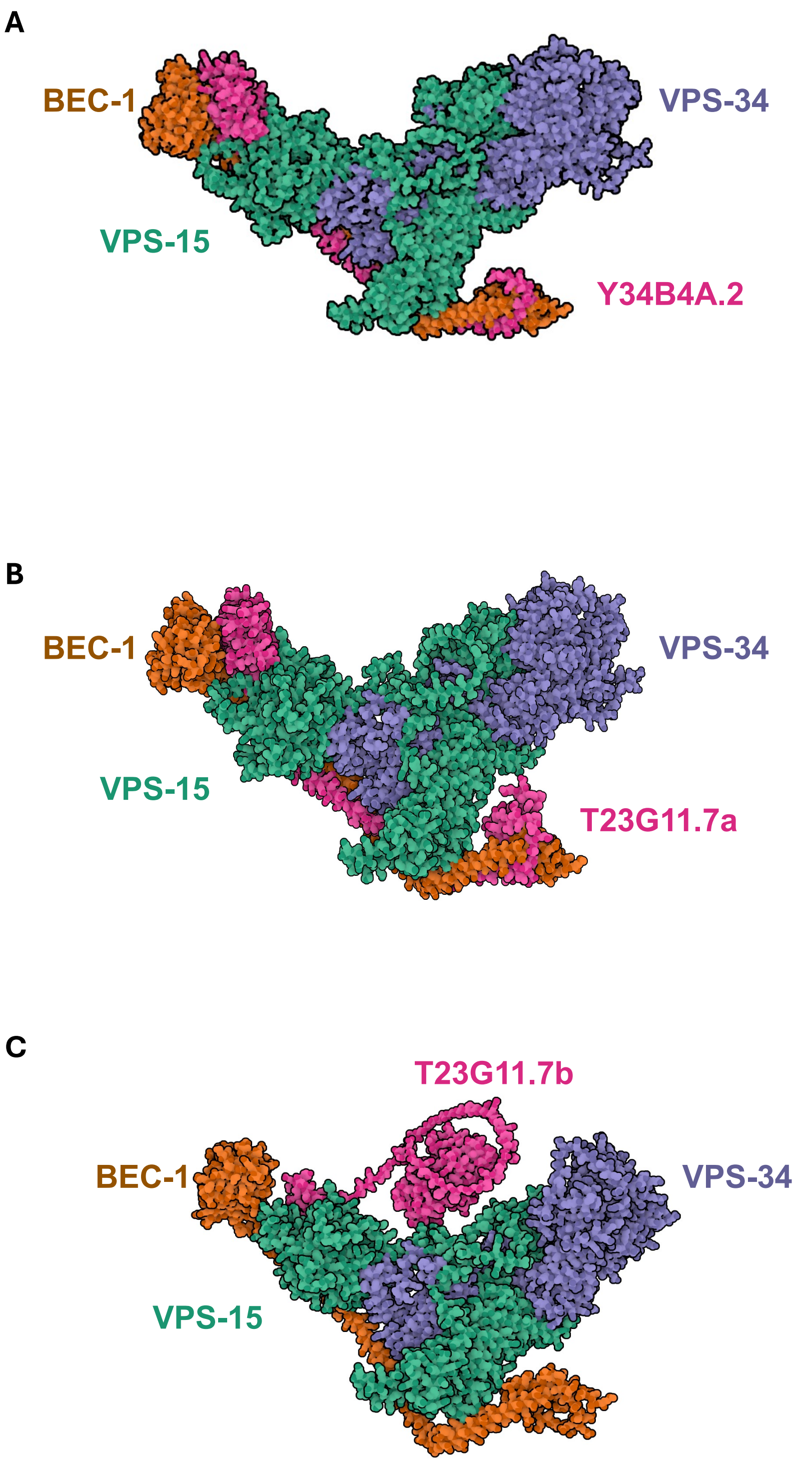


**Supplemental Figure 1:** Structural predictions of *C. elegans* UVRAG orthologs in complex with VPS-34, VPS-15, and BEC-1. **(A)** Y34B4A.2 predicted structural conformation in complex with BEC-1, VPS-15, and VPS-34. **(B)** T23G11.7a predicted structural conformation in complex with BEC-1, VPS-15, and VPS-34 **(C)** T23G11.7.b predicted structural conformation in complex with BEC-1, VPS-15, and VPS-34.

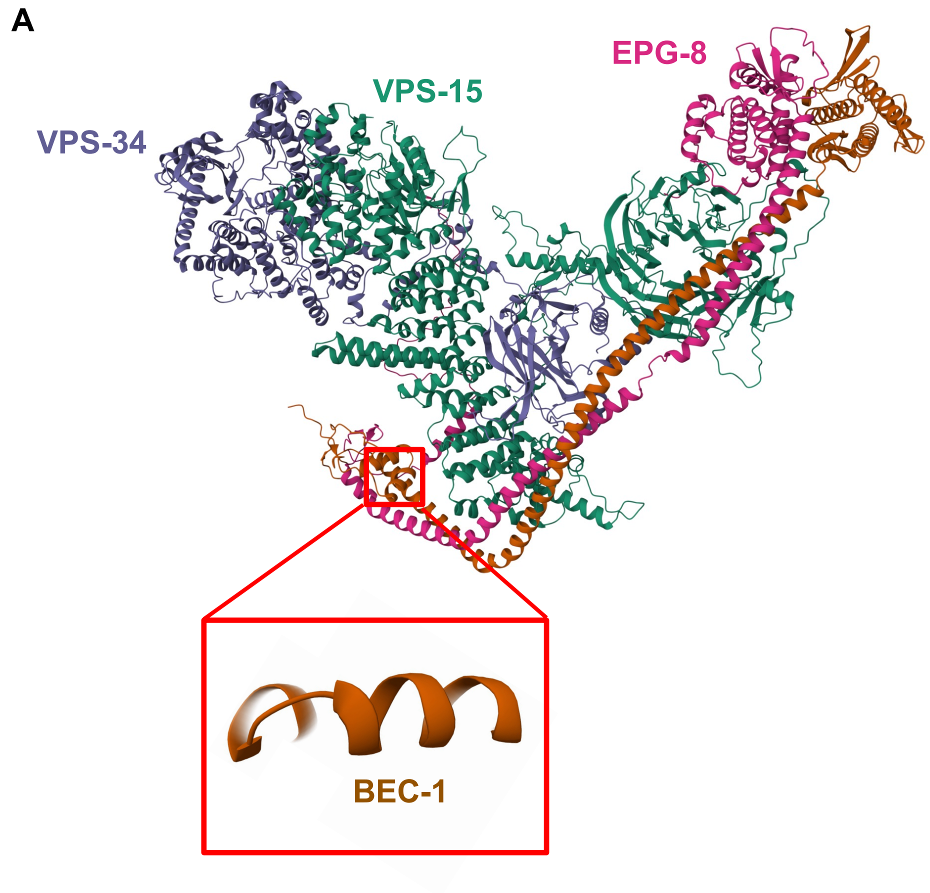


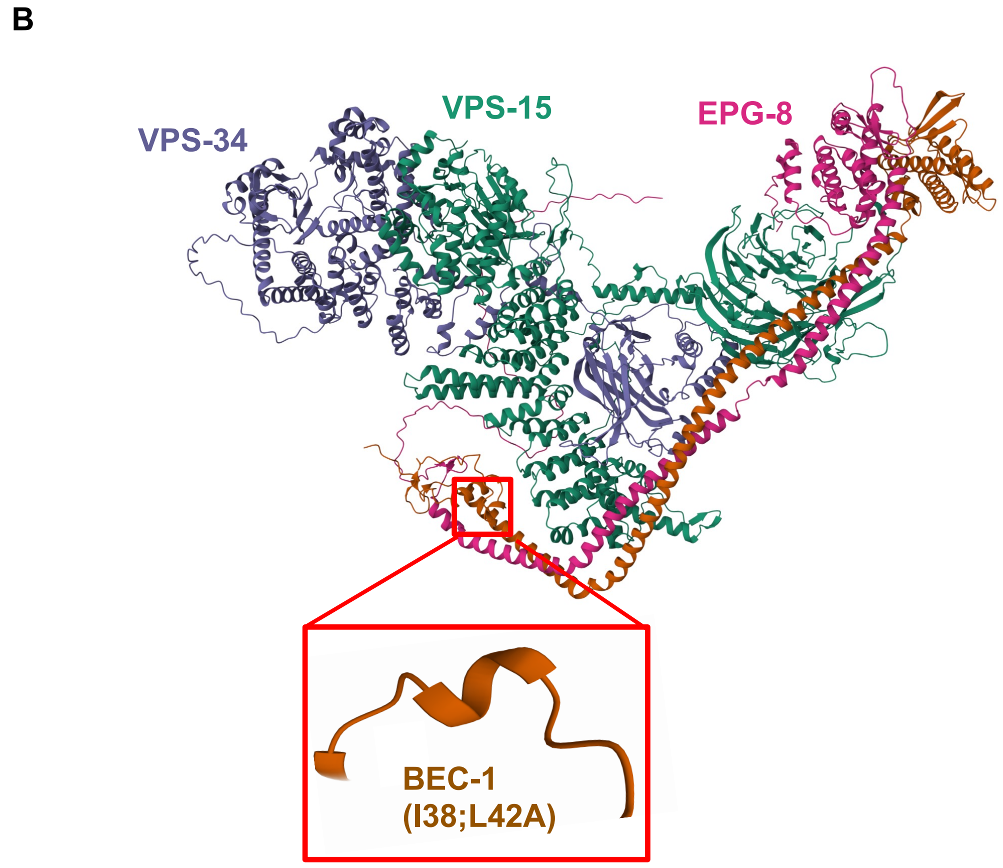


**Supplemental Figure 2. Structural predictions of WT BEC-1 and BEC-1 with mutations in the BH3 domain. (A)** WT BEC-1 in complex with VPS-15, EPG-8, and VPS-34. BH3 domain structure is shown in enlarged inset. **(B)** BEC-1, with residues within the BH3 domain (I38;L42A) domain mutated, in complex with VPS-15, EPG-8, and VPS-34. BH3 domain structure is shown in enlarged inset.
